# CROSS-PLATFORM CONCORDANCE ANALYSIS REVEALS ROBUST AND CLINICALLY RELEVANT GENE SIGNATURES IN BREAST CANCER

**DOI:** 10.64898/2026.10.13.758788

**Authors:** J.O Imhanbor, A.A Fatai

## Abstract

**Background:** Breast cancer is a molecularly heterogeneous disease characterized by diverse transcriptional programs across tumor subtypes. RNA sequencing (RNA-seq) has become a central tool for profiling these alterations; however, variability in data processing pipelines can introduce inconsistencies in downstream analyses, limiting the identification of robust and reproducible gene signatures.

**Methods:** We implemented a three-phase integrative framework to identify high-confidence gene signatures in breast cancer. The framework comprised: (i) a discovery phase involving differential expression analysis across two independent RNA-seq workflows — a STAR-counts pipeline applied to a balanced TCGA-BRCA cohort (30 tumor vs. 30 normal samples) processed via TCGAbiolinks, and a Salmon-based pseudo-alignment pipeline applied to an independent SRA-derived dataset (3 tumor vs. 3 normal samples) processed from raw sequencing reads; (ii) a cross-dataset concordance phase identifying consensus upregulated genes through gene symbol intersection analysis; and (iii) a validation and clinical characterisation phase comprising large-scale validation in the full TCGA-BRCA cohort (n□=□1,093 tumors, 112 normal samples), functional enrichment, PAM50 subtype stratification, survival analysis, and cancer cell line expression profiling. A multi-criteria scoring framework was applied to prioritize potential drug targets.

**Results:** A total of 666 genes were consistently upregulated across both discovery datasets, indicating strong cross-platform concordance. Of these, 502 genes (93%) were validated as upregulated in the full TCGA cohort, demonstrating high reproducibility. Functional enrichment analyses revealed dominant involvement in cell cycle regulation, chromosomal segregation, and DNA repair pathways. Subtype analysis showed a marked enrichment of these genes in Basal-like and HER2-enriched tumors, highlighting their association with more aggressive disease phenotypes. Survival analysis identified a subset of genes significantly associated with poor overall survival, while cell line validation confirmed expression in breast cancer models. Integration of these features enabled the prioritization of candidate genes with potential therapeutic relevance.

**Conclusion:** This study demonstrates that integrating two independent RNA-seq analytical workflows (a STAR-counts pipeline applied to a balanced TCGA-BRCA discovery cohort and a Salmon-based pipeline applied to an independent SRA-derived dataset) enables the identification of robust and clinically meaningful gene signatures in breast cancer. The high concordance observed across these independent datasets underscores the reproducibility of the identified signals, while downstream analyses highlight their biological and translational relevance. This framework provides a systematic approach for prioritizing candidate biomarkers and therapeutic targets in cancer transcriptomics.

## Introduction

Breast cancer remains one of the most extensively studied malignancies worldwide and is characterized by substantial molecular and clinical heterogeneity. Advances in large-scale transcriptomic profiling have revealed that breast cancer comprises biologically distinct subtypes, including Luminal A, Luminal B, HER2-enriched, and Basal-like tumors, each associated with unique gene expression programs, therapeutic responses, and clinical outcomes (Perou et al., 2000; Parker et al., 2009; Cancer Genome Atlas Network, 2012). These discoveries have transformed transcriptomics into a central component of breast cancer research, enabling improved molecular classification, biomarker discovery, and therapeutic stratification.

The widespread adoption of RNA sequencing (RNA-seq) has significantly accelerated the characterization of tumor-associated transcriptional alterations by providing high-resolution, genome-wide expression profiling (Wang et al., 2009). More recently, transcriptomic studies have increasingly integrated large patient cohorts, multi-omics datasets, and computational approaches to identify clinically actionable molecular signatures in breast cancer (Curtis et al., 2012; Jiang et al., 2022). In parallel, the emergence of lightweight transcript quantification tools such as Salmon and advanced statistical frameworks such as DESeq2 has improved the efficiency and scalability of RNA-seq analysis pipelines (Patro et al., 2017). However, despite these methodological advances, reproducibility remains a major challenge in computational transcriptomics.

Differences in alignment strategies, transcript quantification methods, normalization procedures, and filtering thresholds can substantially influence downstream differential expression results, leading to inconsistencies in identified gene signatures across studies (Conesa et al., 2016; Fonseca et al., 2024). Recent benchmarking efforts have demonstrated that such variations in computational workflows can alter biological interpretation, pathway enrichment outcomes, and candidate biomarker prioritization, particularly in heterogeneous cancers such as breast cancer (Merino et al., 2019). At the same time, the growing emphasis on precision oncology has highlighted the need for transcriptomic signatures that are not only statistically significant but also biologically robust and clinically reproducible across independent cohorts and analytical pipelines (Lähnemann et al., 2020; Zhang et al., 2023). Despite these concerns, only few studies have systematically assessed whether tumor-associated transcriptional signatures remain concordant across independently processed RNA-seq workflows in breast cancer, nor whether signatures identified through cross-platform concordance retain biological and clinical relevance at population scale.

Molecular subtype classification systems such as PAM50 have demonstrated that aggressive subtypes, particularly Basal-like and HER2-enriched tumors, are associated with elevated proliferative activity, genomic instability, and altered immune signaling pathways (Parker et al., 2009; Cancer Genome Atlas Network, 2012). Recent transcriptomic and single-cell studies have further shown that these subtypes exhibit extensive intra-tumoral heterogeneity and dynamic pathway activation patterns linked to disease progression and treatment resistance (Wu et al., 2021; Kim et al., 2024). Consequently, identifying gene signatures that remain consistent across datasets and analytical strategies may improve the reliability of candidate biomarkers and facilitate the prioritization of therapeutically relevant targets.

In this study, we developed a three-phase integrative framework to identify reproducible and clinically relevant gene signatures in breast cancer through cross-platform transcriptomic validation. The framework comprised: (i) a discovery phase in which differential expression analysis was independently performed using a STAR-counts-based pipeline applied to a balanced TCGA-BRCA cohort (30 tumor vs. 30 normal samples) and a Salmon-based pseudo-alignment pipeline applied to an independent SRA-derived dataset (3 tumor vs. 3 normal samples) processed from raw sequencing reads; (ii) a cross-dataset concordance phase in which consensus upregulated genes were identified through gene symbol intersection analysis across both discovery datasets; and (iii) a validation and clinical characterisation phase encompassing large-scale validation in the full TCGA-BRCA cohort, functional enrichment, PAM50 subtype stratification, survival analysis, and breast cancer cell line expression profiling.

To assess the broader biological and clinical significance of these findings, we validated the consensus gene set in the full TCGA-BRCA cohort comprising over one thousand tumor samples and conducted downstream functional enrichment, PAM50 subtype stratification, survival analysis, and breast cancer cell line validation. We further implemented a multi-criteria evidence scoring framework to prioritize candidate genes with potential therapeutic relevance.

By integrating two independent RNA-seq analytical workflows with large-scale validation and clinically oriented downstream analyses, this study addresses a critical gap in cancer transcriptomics: the systematic identification of molecular signatures that are concordant across computational pipelines while retaining biological and translational significance. Our findings provide a reproducible framework for RNA-seq-based biomarker discovery and contribute to ongoing efforts in precision oncology and therapeutic target prioritization in breast cancer.

## Materials and Methods

### Study Design and Analytical Framework

This study employed a three-phase integrative transcriptomic framework to identify reproducible and clinically relevant gene signatures in breast cancer through cross-platform validation and downstream biological characterization. The framework comprised: (i) a discovery phase involving independent differential expression analysis using two distinct RNA-seq analytical strategies — a STAR-counts pipeline applied to a balanced TCGA-BRCA cohort, and a Salmon-based pseudo-alignment pipeline applied to an independent SRA-derived dataset; (ii) a cross-dataset concordance phase in which consensus upregulated genes were identified through gene symbol intersection analysis; and (iii) a validation and clinical characterisation phase consisting of large-scale validation in the full TCGA-BRCA cohort, functional enrichment, molecular subtype characterization, survival analysis, breast cancer cell line validation, and evidence-based therapeutic target prioritization.

The analytical strategy was designed to evaluate whether tumor-associated transcriptional signals remained consistent across independent datasets and RNA-seq processing workflows while retaining biological and clinical relevance within a large patient cohort. Rather than applying computational batch correction across datasets, the cross-dataset concordance design was used as a conservative filter, retaining only genes consistently upregulated across both datasets despite differences in sequencing origin, quantification methodology, and cohort composition.

### TCGA-BRCA Discovery Cohort

RNA-seq count data and associated clinical metadata for breast invasive carcinoma were obtained from The Cancer Genome Atlas (TCGA-BRCA) using the TCGAbiolinks Bioconductor package (Colaprico et al., 2016). To mitigate the statistical bias inherent in the heavily unbalanced TCGA-BRCA dataset (where tumors vastly outnumber controls), we utilized a class-balanced discovery subset (*n*□=□60) to ensure equal weighting of the variance from normal and malignant tissues during initial signal detection.

Gene expression quantification was based on the Genomic Data Commons (GDC) STAR-counts workflow. Sample annotations were curated to distinguish tumor and normal tissues prior to downstream analysis.

### Independent RNA-seq Dataset Acquisition and Processing

To independently evaluate transcriptomic concordance across analytical pipelines, an additional breast cancer RNA-seq dataset consisting of three primary tumor and three adjacent normal breast tissue samples was analyzed from raw sequencing reads. Sequencing data were obtained from the NCBI Sequence Read Archive using the SRA Toolkit (v3.0) within a Windows Subsystem for Linux (WSL-Ubuntu) environment.

Raw sequencing reads were downloaded using prefetch, followed by FASTQ generation using fasterq-dump with paired-end splitting enabled. Transcript quantification was subsequently performed using Salmon v1.10.3 (Patro et al., 2017). Reference transcript sequences and gene annotations were obtained from GENCODE release v46. A transcriptome index was generated using a k-mer length of 31, after which transcript abundance estimation was performed in quasi-mapping mode with automatic library type detection.

Transcript-level abundance estimates generated for each sample were imported into R and summarized to gene-level counts using the tximport package (Soneson et al., 2015). Transcript-to-gene mapping was derived from the GENCODE v46 annotation file using GenomicFeatures.

### Differential Expression Analysis

Differential expression analysis for both discovery datasets was performed using the DESeq2 package (Love et al., 2014). Prior to analysis, lowly expressed genes were filtered to reduce statistical noise and improve dispersion estimation. Differential expression modeling was conducted using a design formula based on tissue condition (tumor versus normal).

Variance stabilizing transformation (VST) was applied for exploratory analyses and visualization, including principal component analysis (PCA) and hierarchical clustering. To improve effect size estimation for low-count genes, log2 fold-change shrinkage was performed using the apeglm method (Zhu et al., 2019).

Genes exhibiting an adjusted p-value below 0.05 and an absolute log□fold-change greater than 1 were considered significantly differentially expressed following Benjamini–Hochberg multiple testing correction.

### Cross-Dataset Concordance Analysis

To identify reproducible tumor-associated transcriptional signatures, significantly upregulated genes from both discovery datasets were independently standardized and filtered prior to overlap analysis. Ensembl gene identifiers were harmonized by removing version suffixes, and duplicated gene symbols were resolved by retaining entries with the lowest adjusted p-value.

Consensus upregulated genes were subsequently identified through gene symbol intersection analysis in R. This approach enabled the identification of genes consistently upregulated across independent datasets processed using distinct RNA-seq analytical workflows. This concordance-based design served as a conservative batch-robust filter: only genes upregulated in both datasets, despite differences in cohort composition, library preparation, and quantification methodology were retained for downstream analysis.

### Functional Enrichment and Pathway Analysis

Functional enrichment analyses were conducted to characterize the biological processes and pathways associated with the consensus upregulated gene set. Gene Ontology (GO), Kyoto Encyclopedia of Genes and Genomes (KEGG), and Reactome pathway enrichment analyses were performed using the clusterProfiler (Yu et al., 2012) and ReactomePA (Yu & He, 2016) packages.

Gene identifiers were converted from gene symbols to Entrez identifiers using the bitr() function and the org.Hs.eg.db (Carlson, 2019) annotation database. Over-representation analyses were performed across Gene Ontology Biological Process, Molecular Function, and Cellular Component categories. Pathways with adjusted p-values below 0.05 were considered significantly enriched.

Gene Set Enrichment Analysis (GSEA) was additionally performed using ranked gene lists generated from shrunken log□fold-change values. Duplicate Entrez identifiers were resolved through aggregation, and minimal random noise was introduced where necessary to eliminate ranking ties during enrichment computation.

### Validation in the Full TCGA-BRCA Cohort

Consensus upregulated genes identified through cross-dataset concordance analysis were validated using the full TCGA-BRCA cohort retrieved through the curatedTCGAData Bioconductor package (Ramos et al., 2018). The validation dataset consisted of 1,093 tumor samples and 112 normal breast tissue samples with associated clinical annotations.

Tumor and normal samples were identified using TCGA barcode sample type annotations, where “01” represented primary tumors and “11” represented solid tissue normal samples. Mean expression values were computed separately for tumor and normal groups, and validation log□fold-changes were calculated as:

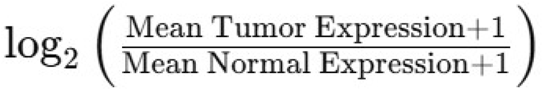

Of the 666 consensus upregulated genes identified, 540 genes were successfully mapped within the full TCGA-BRCA cohort using this approach; the remaining 126 genes were absent from the curatedTCGAData gene annotation set and were therefore excluded from validation analysis. Genes with validation log□fold-change values greater than 1 were considered confirmed upregulated genes within the full TCGA cohort.

### Molecular Subtype Characterization

To investigate subtype-specific transcriptional patterns, PAM50 molecular subtype annotations were extracted from the TCGA-BRCA MultiAssayExperiment object using the PAM50.mRNA metadata field. Subtype-specific mean expression values were computed for each validated gene across Luminal A, Luminal B, HER2-enriched, Basal-like, and Normal-like breast cancer subtypes. For each gene, the dominant subtype association was determined based on the subtype exhibiting the highest mean expression value.

### Survival Analysis

The clinical relevance of validated genes was assessed through overall survival analysis using TCGA clinical metadata. Survival time was defined using days_to_death for deceased patients and days_to_last_followup for censored individuals.

Kaplan–Meier survival analyses were performed using median expression stratification to separate patients into high- and low-expression groups. In addition, univariate Cox proportional hazards regression analyses were conducted using continuous gene expression values to evaluate associations between gene expression and overall survival outcomes. Hazard ratios (HR) and 95% confidence intervals (CI) were reported for each gene.

Multiple testing correction was performed using the Benjamini–Hochberg method, and genes with adjusted p-values below 0.05 were considered significantly associated with survival.

### Breast Cancer Cell Line Validation

To evaluate experimental tractability and translational relevance, expression of validated genes was assessed in breast cancer cell lines obtained from the Cancer Cell Line Encyclopedia (CCLE) through the depmap Bioconductor package. TPM-normalized RNA expression matrices and associated metadata from the 22Q2 release were utilized for analysis.

Cell lines annotated under the “Breast Cancer” primary disease category were retained, and expression profiles of validated genes were examined across the resulting breast cancer cell line panel.

### Evidence-Based Target Prioritization

An evidence-based scoring framework was implemented to prioritize candidate therapeutic targets among the validated genes. Each gene was assigned a composite evidence score (maximum 10 points) based on the following criteria and weightings: differential expression strength across discovery datasets (0–3 points, based on mean log□FC across both cohorts); validation strength in the full TCGA cohort (0–2 points, based on validation log□FC); subtype specificity (0–1 point, awarded for dominant enrichment in Basal-like or HER2-enriched subtypes); survival association (0–2 points, awarded for significant Kaplan–Meier stratification and/or significant Cox regression after multiple testing correction); and expression in breast cancer cell lines (0–1 point, for detectable TPM expression in ≥50% of cell lines).

Additional weighting was assigned to genes with known oncogenic relevance or established druggability, while penalty scores were applied to genes predominantly associated with stromal or extracellular matrix-related processes in order to prioritize tumor-intrinsic candidates. Genes with the highest composite evidence scores were selected as priority targets for downstream therapeutic investigation and future structural or drug discovery analyses.

## Results

### Differential Expression Analysis Reveals Distinct Tumor-Associated Transcriptomic Profiles

Differential expression analysis was initially performed using a controlled TCGA-BRCA discovery cohort consisting of 30 primary tumor samples and 30 normal breast tissue samples. Following count filtering and normalization, variance-stabilizing transformation demonstrated clear separation between tumor and normal samples, indicating strong transcriptomic divergence associated with malignant transformation. Principal component analysis (PCA) revealed that tumor status represented the dominant source of transcriptional variation within the dataset, with tumor and normal samples clustering distinctly across the first principal component.

Differential expression analysis identified thousands of significantly dysregulated genes between tumor and normal tissues using an adjusted p-value threshold of 0.05 and an absolute log□fold-change threshold greater than 1. Volcano plot visualization demonstrated widespread transcriptional remodeling (Figure 1), with both strongly upregulated and downregulated gene populations observed in tumor samples. Hierarchical clustering using the top differentially expressed genes showed robust segregation of tumor and normal samples, further supporting the presence of a strong biological signal within the dataset (Figure 2).

**Figure 1.**
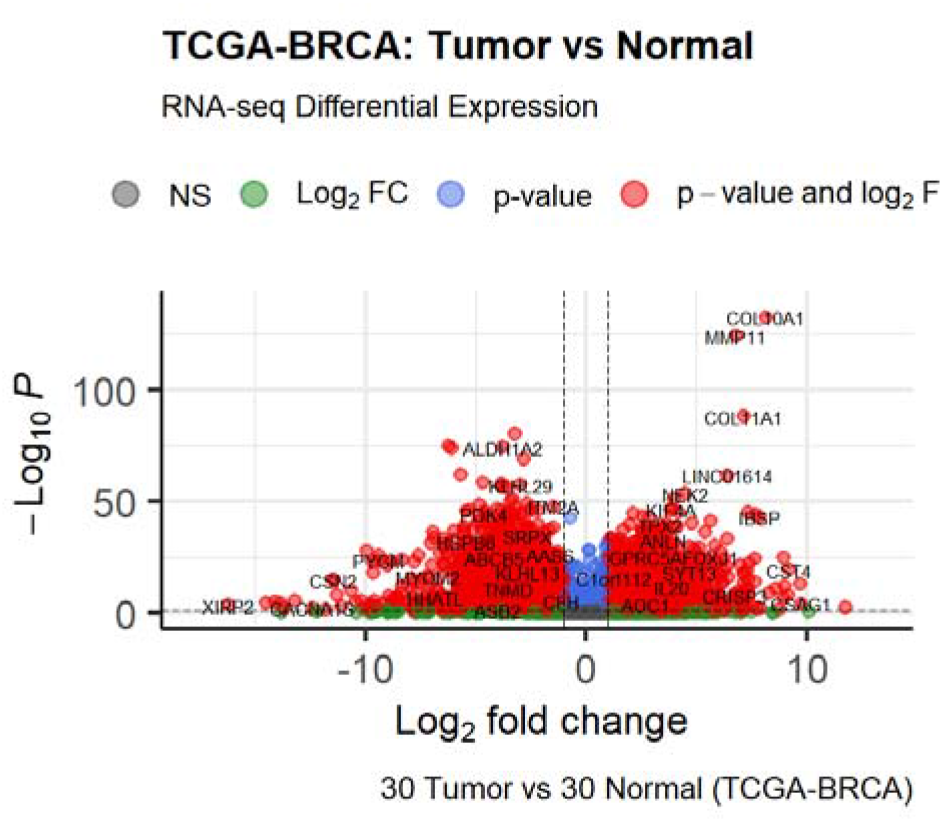
Volcano plot of differential expression analysis for the TCGA 30 vs. 30 discovery cohort. Each point represents a gene; the x-axis shows log□ fold-change (tumor vs. normal) and the y-axis shows −log□ □adjusted p-value. Red points indicate significantly upregulated genes (log□FC > 1, adj. p < 0.); blue points indicate significantly downregulated genes. Dashed lines represent the significance thresholds applied.

**Figure 2.**
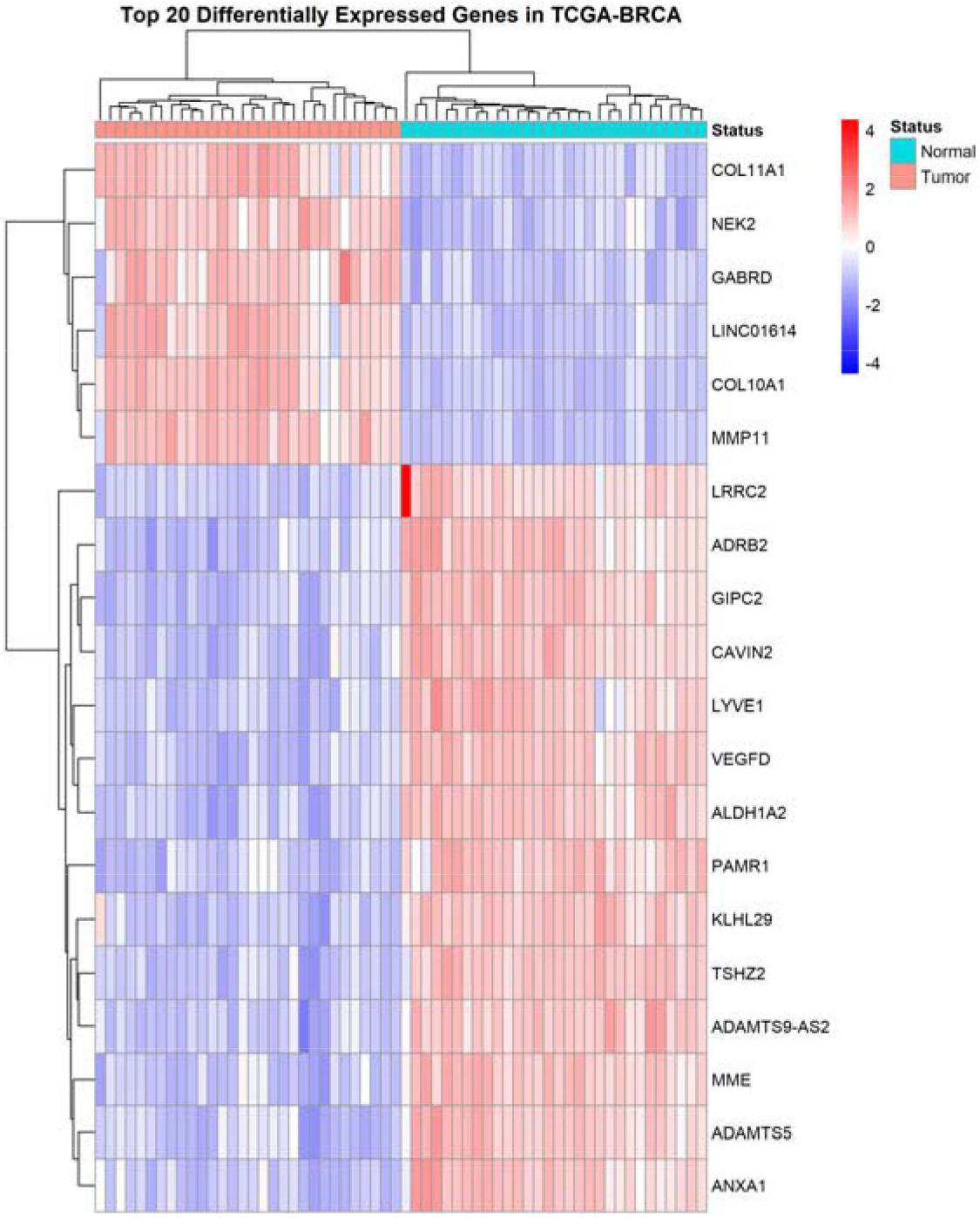
Heatmap of the top 20 differentially expressed genes in the TCGA 30 vs. 30 discovery cohort. Rows represent genes; columns represent samples. Color intensity indicates variance-stabilized expression values. Tumor and normal samples are annotated by column color bar.

To independently assess transcriptomic reproducibility across analytical workflows, a second breast cancer RNA-seq dataset consisting of three tumor and three adjacent normal tissue samples was processed from raw sequencing reads using a Salmon-based quantification pipeline. PCA analysis similarly demonstrated strong separation between tumor and adjacent normal tissues, with the first principal component explaining 54% of total variance and the second principal component explaining 22% of variance (Figure 3).

**Figure 3.**
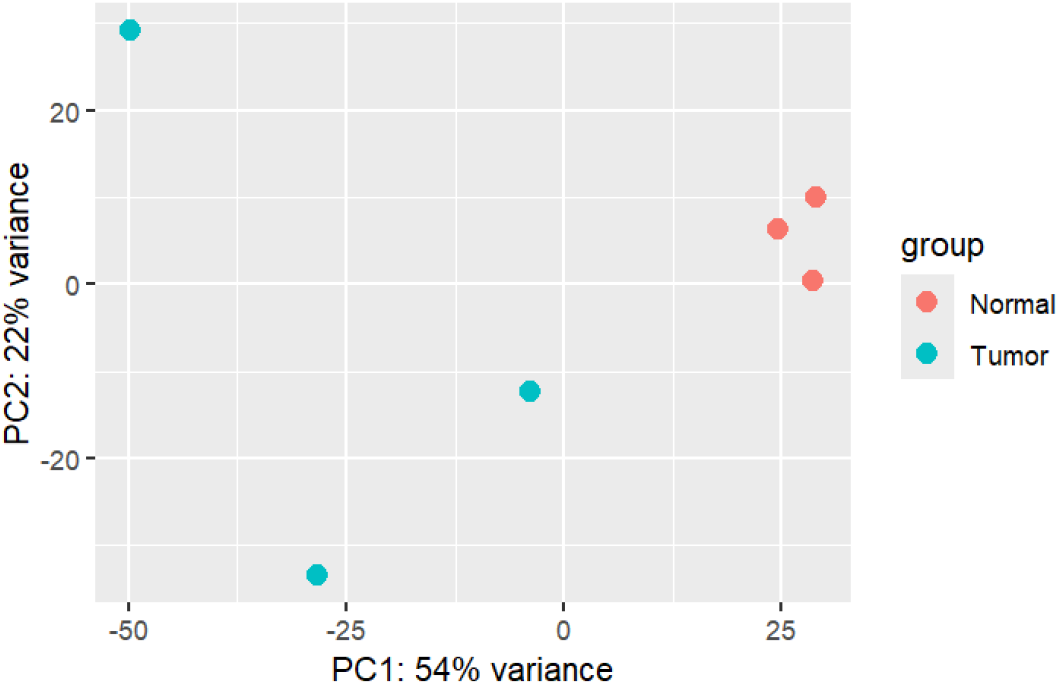
PCA plot of the independent Salmon-processed RNA-seq dataset (3 tumor, 3 normal samples). Each point represents one sample. PC1 explained 54% of total variance and PC2 explained 22% of total variance, with clear separation between tumor and adjacent normal samples.

Differential expression analysis of the independently processed dataset identified 3,008 significantly dysregulated genes, of which 2,640 exhibited absolute log□fold-change values greater than 1. These findings indicated extensive transcriptional reprogramming associated with breast tumor tissue and demonstrated that biologically meaningful expression signals remained detectable despite differences in dataset origin and RNA-seq processing strategy (Figure 4).

**Figure 4.**
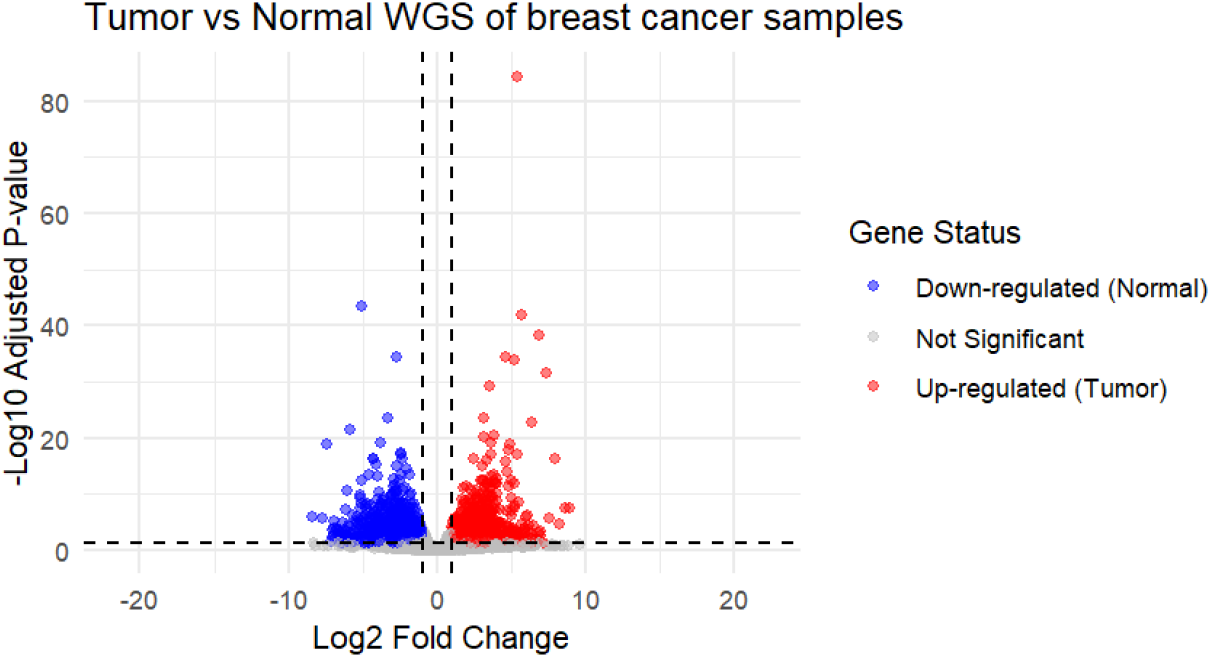
Volcano plot of differential expression analysis for the independent Salmon-processed dataset (3 tumor vs. 3 normal). Axes and color coding as described for Figure 1. A total of 3,008 significantly dysregulated genes were identified, of which 2,640 exhibited |log□FC| > 1.

### Cross-Dataset Concordance Identifies Reproducible Upregulated Gene Signatures

To identify robust tumor-associated transcriptional signatures, significantly upregulated genes from both discovery datasets were compared through cross-dataset concordance analysis. After identifier harmonization, filtering, and duplicate removal, 4,103 upregulated genes were retained from the TCGA 30 vs. 30 cohort, while 1,328 upregulated genes were retained from the independently processed Salmon-based dataset.

Intersection analysis identified 666 genes consistently upregulated across both datasets, demonstrating substantial cross-platform concordance despite differences in cohort composition, sequencing origin, and transcript quantification methodology (Figure 5). The high degree of overlap observed across independent workflows suggested that the identified consensus gene set represented biologically stable transcriptional alterations associated with breast tumorigenesis rather than workflow-specific analytical artifacts.

**Figure 5.**
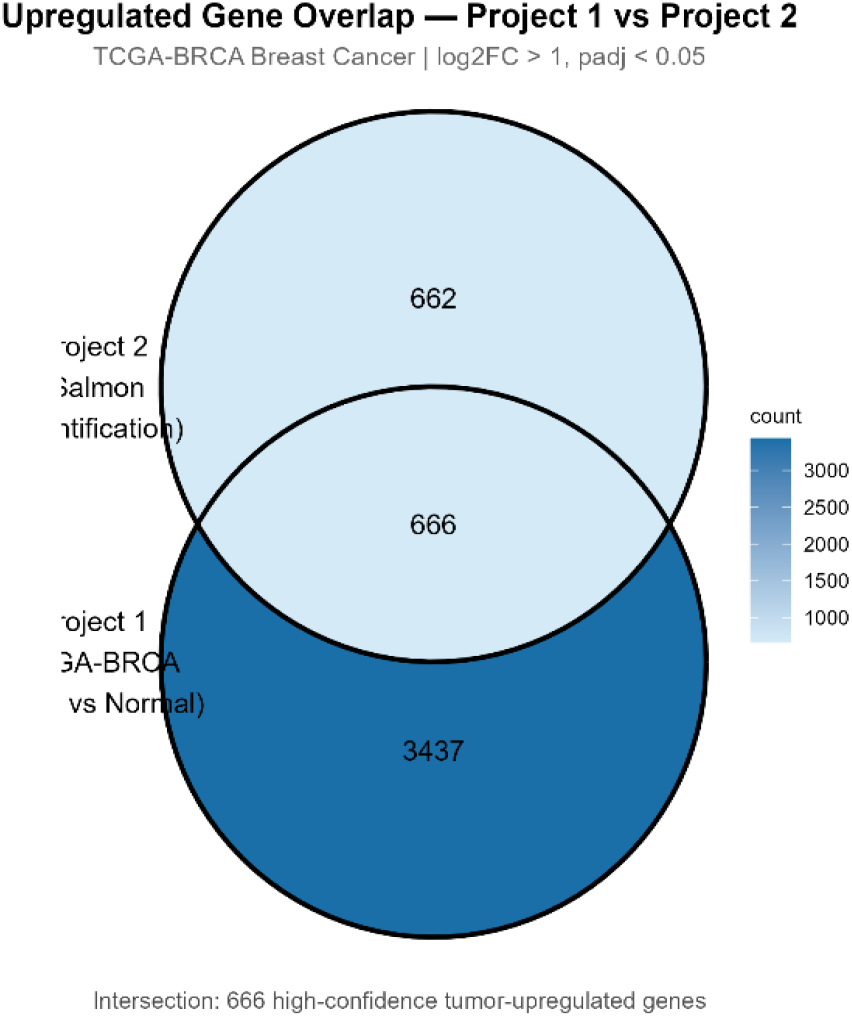
Venn diagram illustrating the overlap of significantly upregulated genes between the two discovery datasets. After harmonization and filtering, 4,103 upregulated genes were identified in the TCGA 30 vs. 30 cohort and 1,328 in the Salmon-processed dataset, yielding 666 consensus upregulated genes present in both.

### Functional Enrichment Analysis Reveals Dominant Cell Cycle and Genomic Instability Signatures

Functional enrichment analyses were performed to characterize the biological significance of the 666 consensus upregulated genes. Gene Ontology Biological Process analysis identified strong enrichment for pathways associated with mitotic progression, chromosome segregation, nuclear division, and DNA replication. Among the most significantly enriched biological processes were mitotic sister chromatid segregation, organelle fission, chromosome segregation, and nuclear division, collectively indicating extensive dysregulation of proliferative and cell cycle-associated programs in breast tumors (Figure 6).

**Figure 6.**
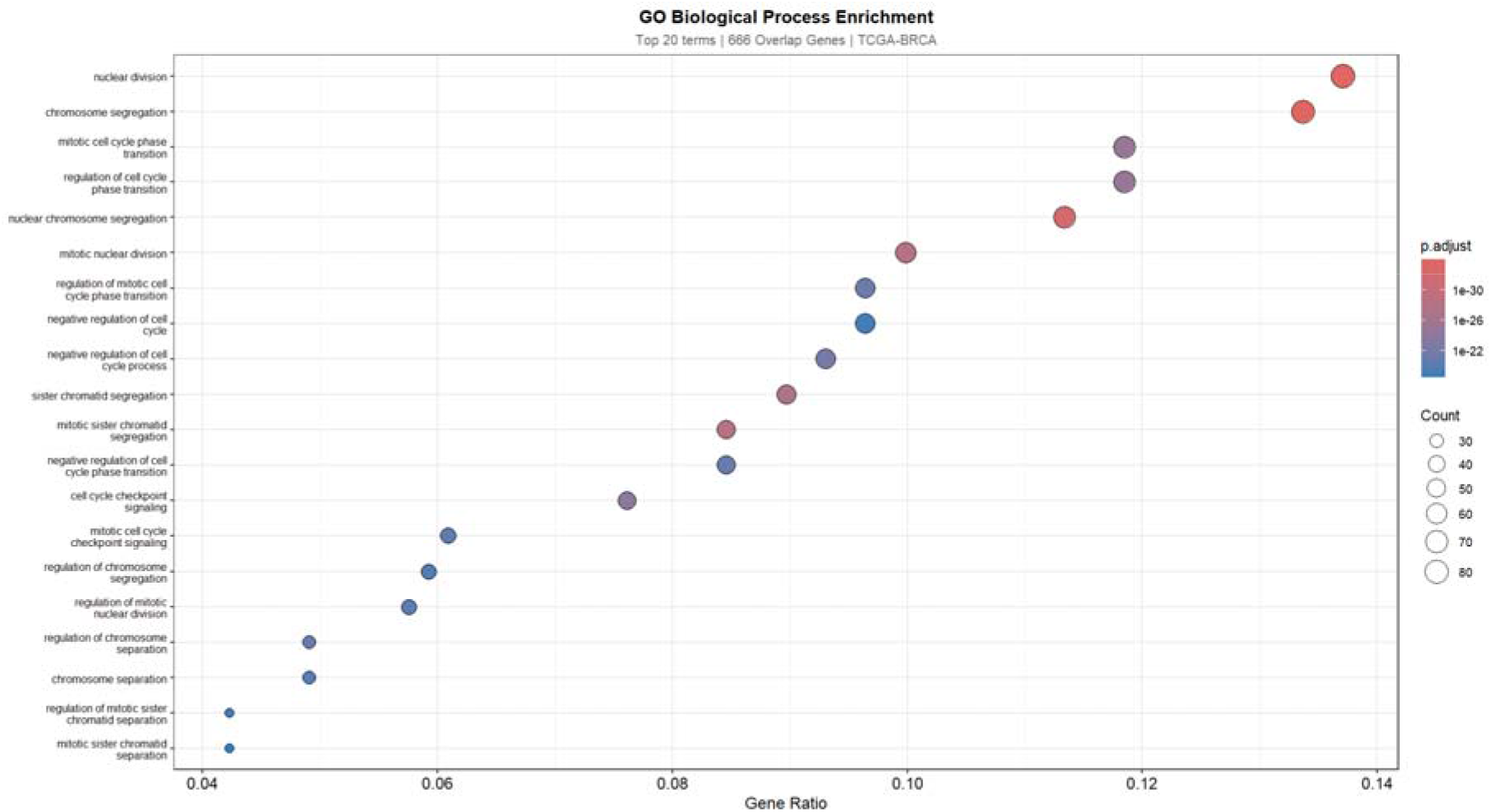
Gene Ontology (GO) Biological Process enrichment dot plot for the 666 consensus upregulated genes. Each dot represents a significantly enriched GO term (adj. p < 0.05). Dot size indicates gene count; dot color represents adjusted p-value. KEGG and Reactome enrichment analyses yielded concordant results and are presented in Supplementary Figures 1 and 2.

**Figure 7.**
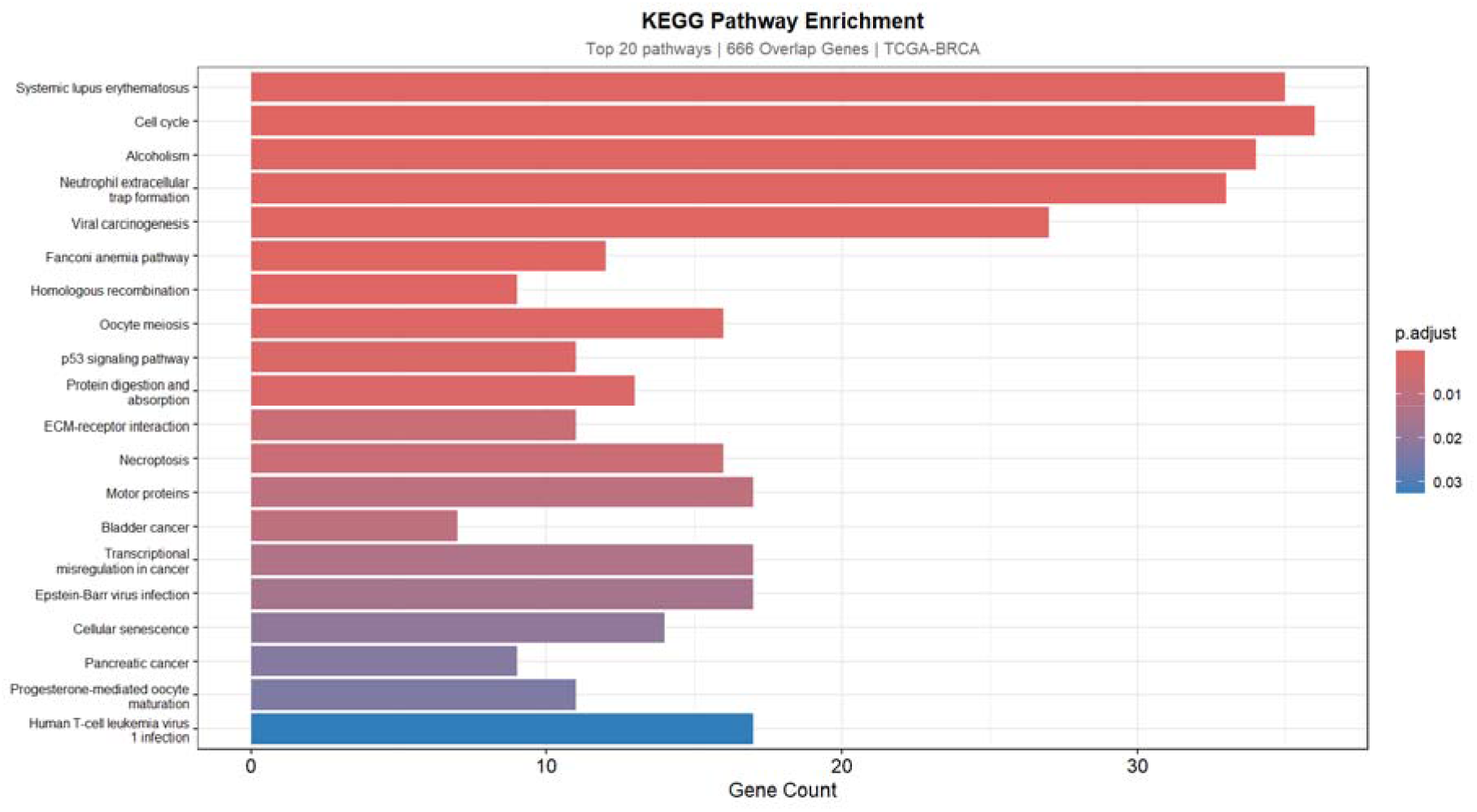
KEGG pathway enrichment plot for the 666 consensus upregulated genes. Significantly enriched pathways (adj. p < 0.05) are shown. Top enriched pathways include cell cycle, Fanconi anemia, homologous recombination, DNA replication, and p53 signaling.

**Figure 8.**
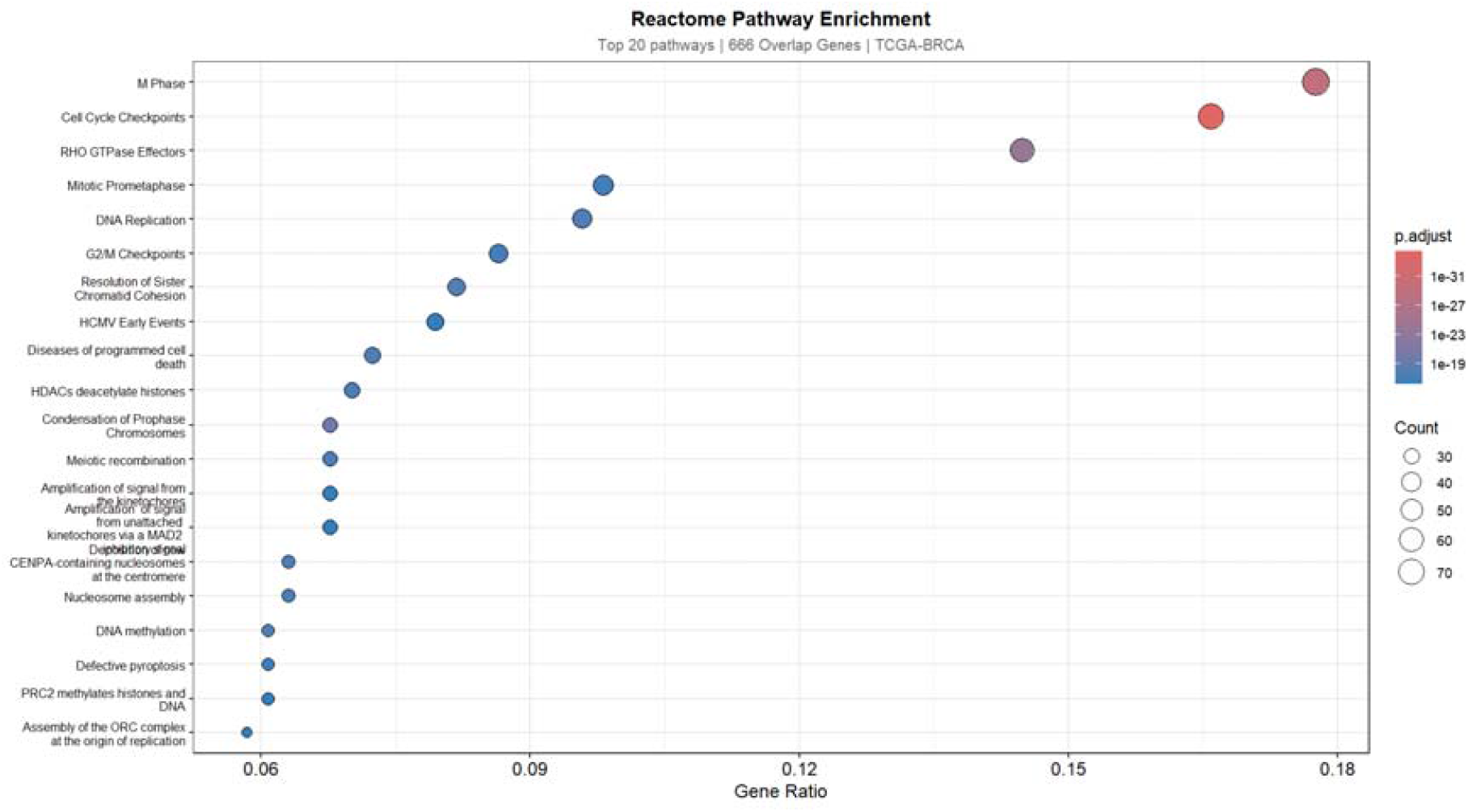
Reactome pathway enrichment analysis for the 666 consensus upregulated genes. Enriched pathway categories include M phase progression, cell cycle checkpoints, DNA damage response, and chromosomal maintenance mechanisms (adj. p < 0.05).

KEGG pathway analysis (Kanehisa & Goto, 2000) further demonstrated significant enrichment of pathways associated with cell cycle regulation, Fanconi anemia signaling, homologous recombination, DNA replication, and p53 signaling. Reactome pathway analysis (Fabregat et al., 2018) reinforced these findings, revealing extensive enrichment of pathways related to M phase progression, cell cycle checkpoints, DNA damage response, and chromosomal maintenance mechanisms. As these three enrichment analyses yielded highly concordant results, KEGG and Reactome outputs are presented in Supplementary Figures 1 and 2; the GO Biological Process analysis is shown as the primary enrichment result (Figure 6). Collectively, these findings indicated that the consensus gene set was strongly associated with proliferative signaling and genomic instability, both of which represent hallmark biological features of aggressive breast cancer phenotypes.

### Validation in the Full TCGA-BRCA Cohort Confirms High Reproducibility of Consensus Genes

To evaluate the reproducibility of the consensus gene signature within a substantially larger patient cohort, validation analysis was performed using the full TCGA-BRCA dataset comprising 1,093 tumor samples and 112 normal breast tissue samples. Among the 666 consensus upregulated genes identified through cross-dataset concordance analysis, 540 genes were successfully mapped within the full TCGA cohort; the remaining 126 genes were absent from the curatedTCGAData gene annotation and were excluded from validation. Of the 540 mapped genes, 502 exhibited validation log□fold-change values greater than 1, corresponding to a validation rate of approximately 93% (502/540).

The high validation rate observed across the independent TCGA cohort demonstrated that the majority of consensus genes retained strong tumor-associated expression patterns at population scale, supporting the robustness and reproducibility of the integrative analytical framework.

### Molecular Subtype Analysis Reveals Enrichment in Aggressive Breast Cancer Subtypes

To investigate subtype-specific transcriptional patterns, expression profiles of the 502 validated genes were examined across PAM50 molecular subtypes within the TCGA-BRCA cohort. Subtype analysis demonstrated that Basal-like tumors exhibited the highest enrichment of validated genes, accounting for 221 genes (44% of the validated signature), followed by HER2-enriched tumors with 113 genes (22%). In contrast, Luminal A tumors exhibited the lowest proportion of dominant subtype-associated genes despite representing the largest patient subgroup (Figure 9).

**Figure 9.**
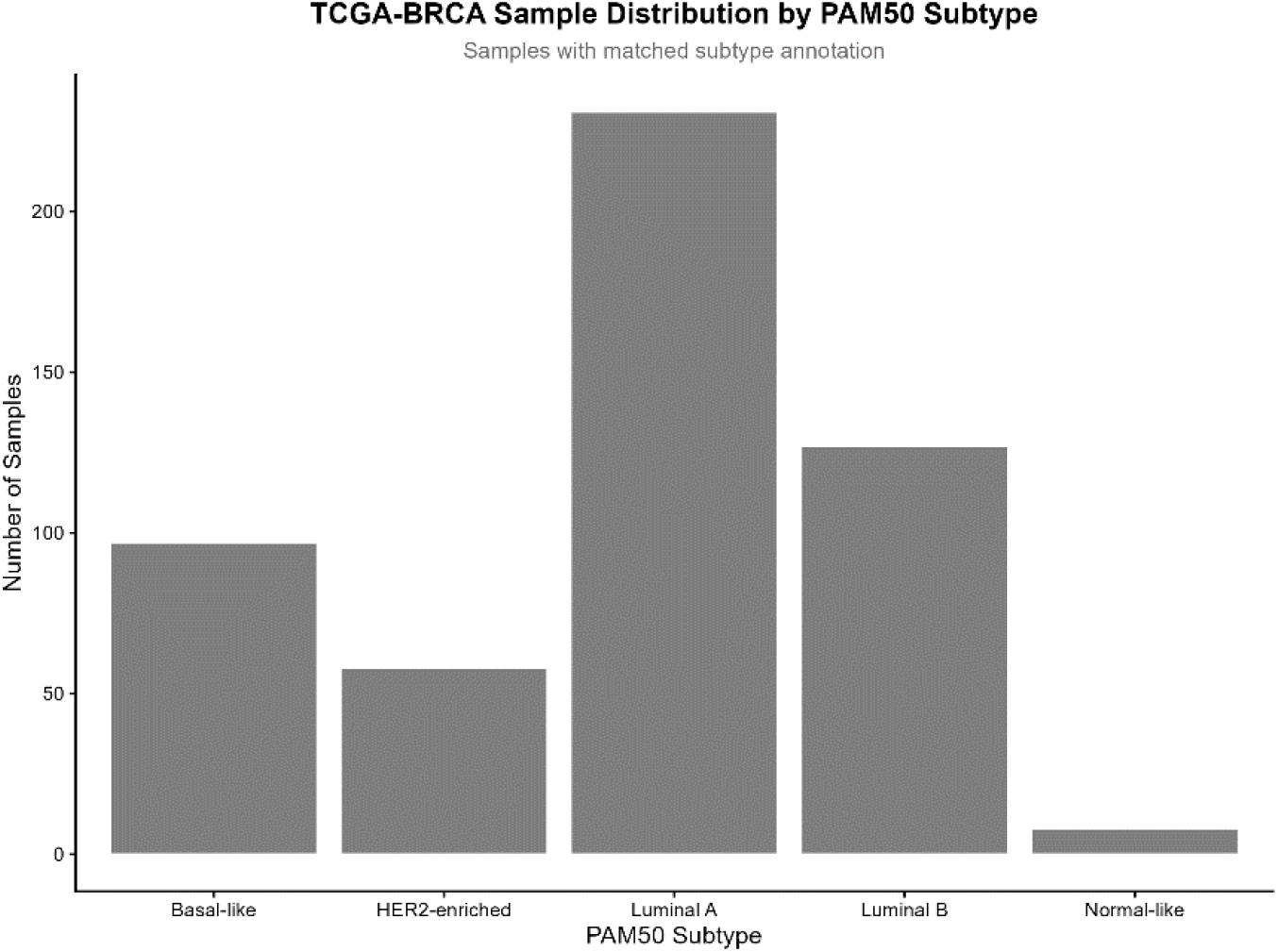
PAM50 subtype distribution bar plot for the 502 validated genes. Bar height indicates the number of validated genes for which that subtype exhibited the highest mean expression. Basal-like tumors account for 22 genes (44%) and HER2-enriched tumors for 113 genes (22%) of the validated signature.

The predominance of validated genes within Basal-like and HER2-enriched tumors suggested that the identified transcriptional signature was strongly associated with biologically aggressive breast cancer phenotypes characterized by elevated proliferative activity and genomic instability. Heatmap visualization further demonstrated subtype-specific expression gradients across validated genes, with pronounced overexpression observed in Basal-like tumors (Figure 10).

**Figure 10.**
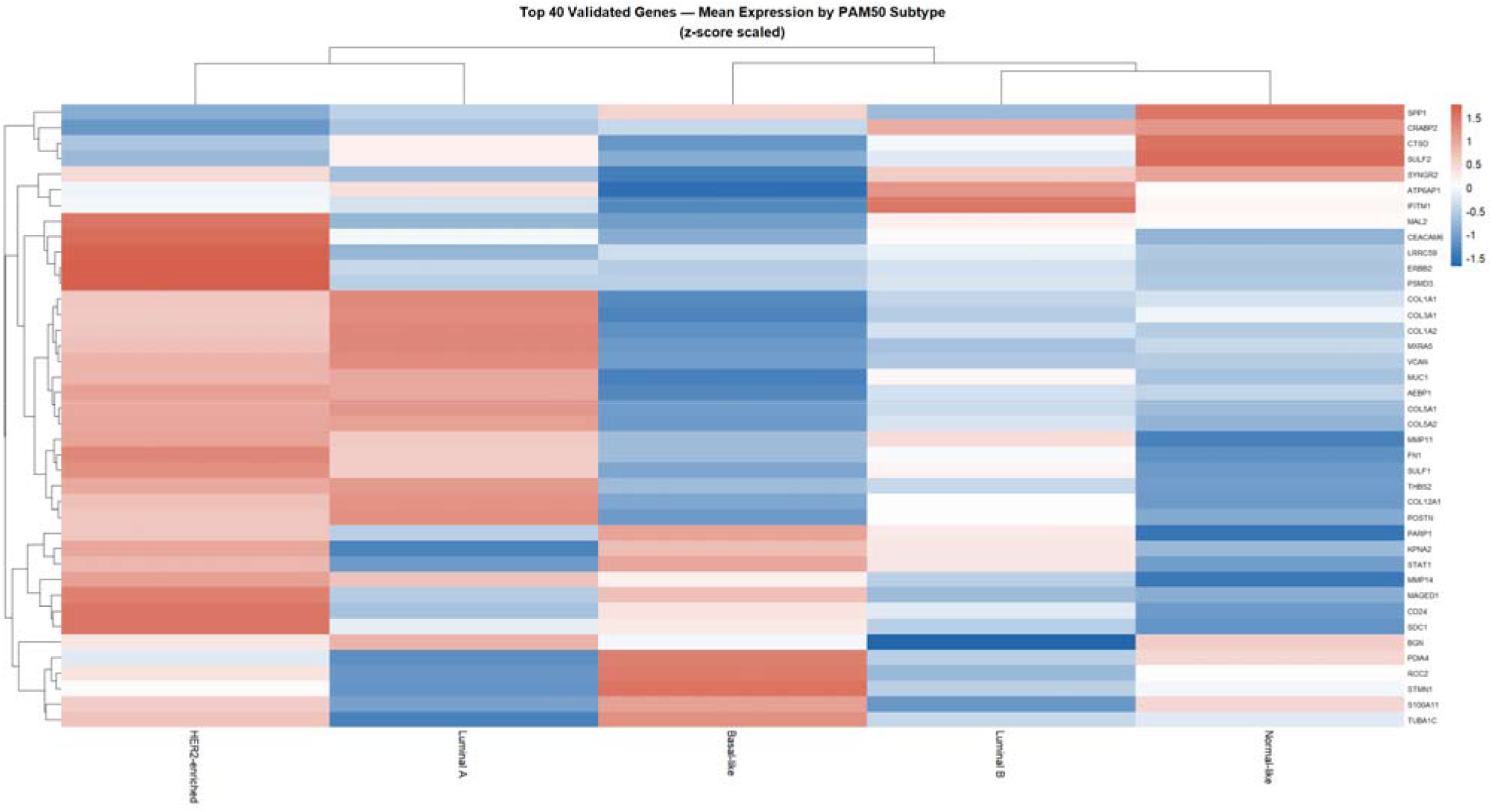
Heatmap of validated gene expression across PAM50 molecular subtypes within the TCGA-BRCA cohort. Rows represent genes; columns represent PAM50 subtypes. Color represents mean VST expression. Pronounced overexpression is observed in Basal-like tumors.

### Survival Analysis Identifies Clinically Relevant Candidate Genes

To assess the clinical relevance of validated genes, survival analyses were conducted using overall survival data from the TCGA-BRCA cohort. Survival time was defined using days_to_death for deceased patients and days_to_last_followup for censored individuals. Due to the relatively low event rate characteristic of TCGA breast cancer cohorts, stringent multiple testing correction considerably narrowed the validated signature to a small subset of genes with robust survival associations.

Univariate Cox proportional hazards regression was performed across all validated genes, followed by Benjamini– Hochberg correction for multiple testing. Of the 502 validated genes tested, four reached statistical significance after correction (adjusted p < 0.05): *MMP13, ULBP2, SHCBP1*, and *MAL2*. Increased expression of each gene was associated with poorer overall survival outcomes, as reflected by hazard ratios greater than 1 in all four cases. The Cox regression results for these genes are summarised in Table 1. While all four genes reached statistical significance following multiple testing correction, effect sizes varied across candidates. *MMP13* demonstrated the most robust association, with the lowest adjusted p-value (padj = 0.000145) and the strongest effect size relative to expression unit. *ULBP2* and *SHCBP1* showed moderate effect sizes with well-bounded confidence intervals, while *MAL2* exhibited a smaller per-unit hazard ratio, reflecting the scale of its expression values. Also several other genes in the validated set, like *SPP1, STMN1*, and *TUBA1C*, did not reach statistical significance in the survival analysis after multiple testing correction (adjusted p > 0.05), indicating that their detection as concordant, validated upregulated genes does not extend to a significant survival association in this cohort.

**Table 1.**
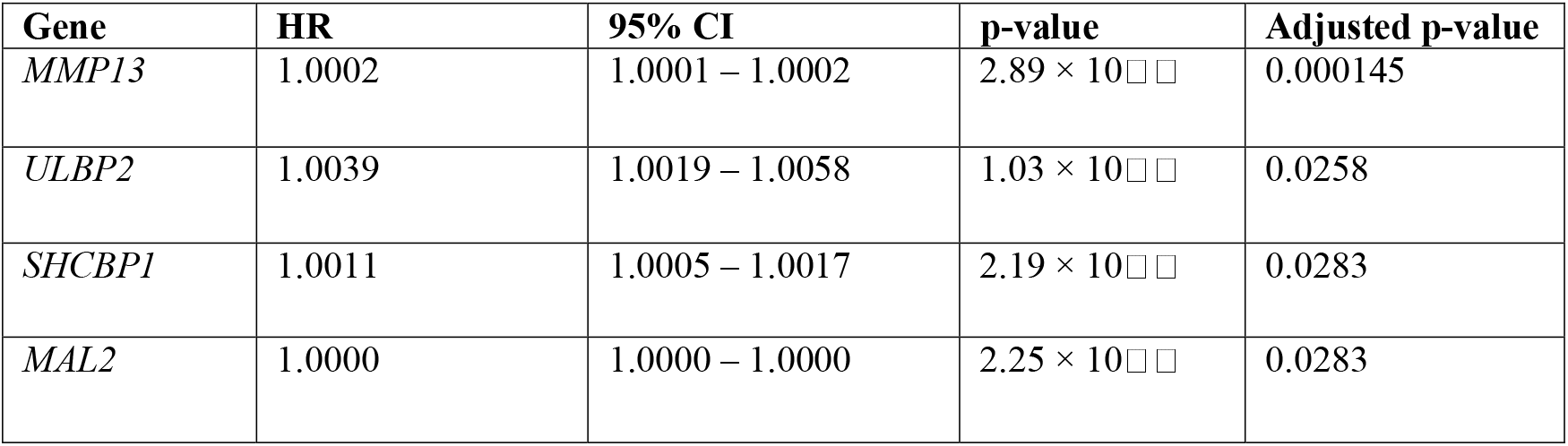
Cox Proportional Hazards Regression Results for Survival-Associated Genes.

### Breast Cancer Cell Line Validation Supports Experimental Tractability

To evaluate the experimental relevance of validated genes, expression profiles were examined across breast cancer cell lines derived from the Cancer Cell Line Encyclopedia (CCLE). A substantial proportion of validated genes demonstrated detectable expression across breast cancer cell lines, indicating that these transcriptional alterations were not restricted to patient tumor tissue and could potentially be investigated in vitro using established experimental models (Figure 11). These findings provided additional support for the translational relevance and experimental tractability of the identified candidate genes.

**Figure 11.**
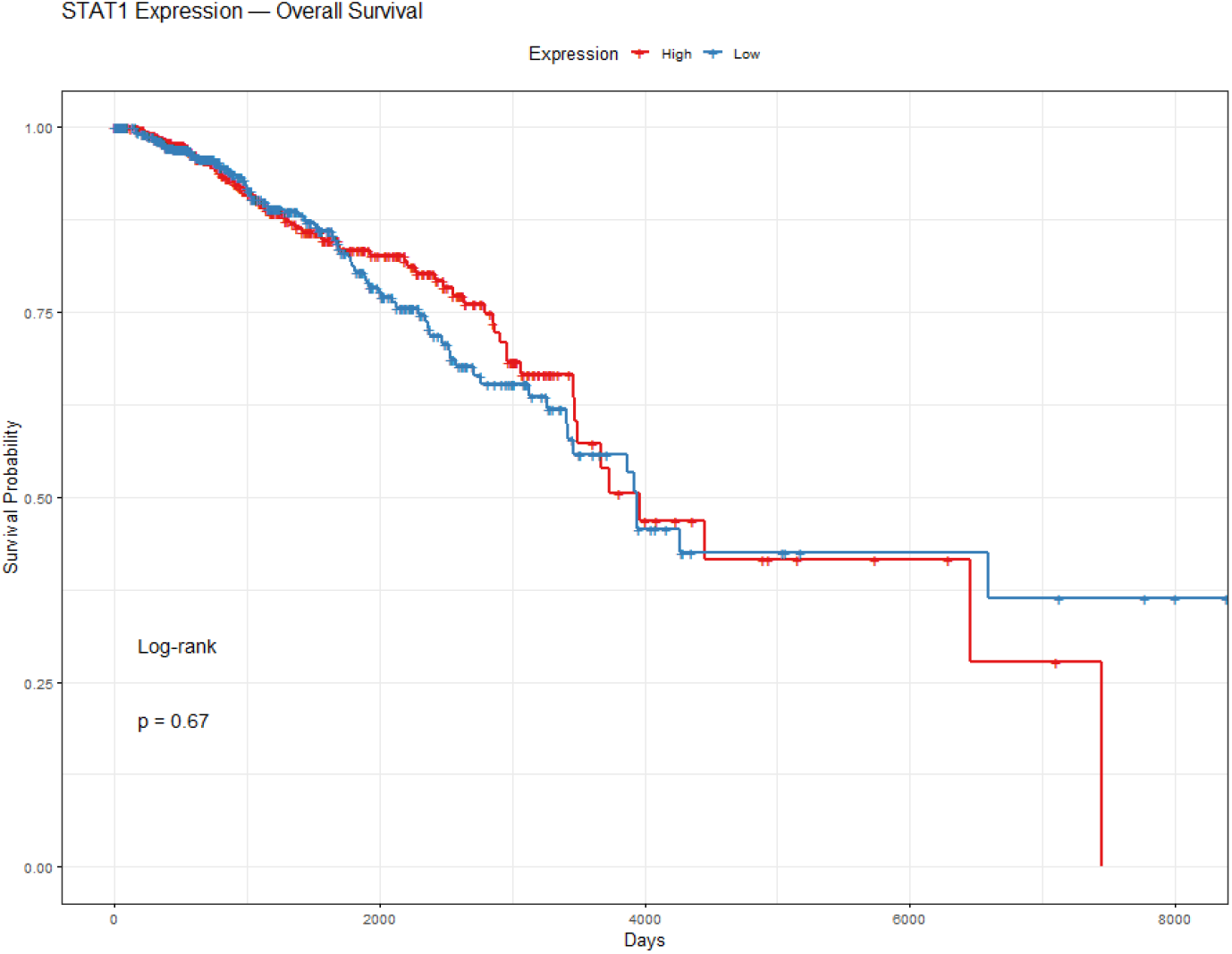
Kaplan–Meier survival curves for STAT1 gene showing survival probability curves when the gene is highly expressed vs lowly expreessed over a period in breast cancer.

### Evidence-Based Prioritization Identifies Candidate Therapeutic Targets

To prioritize candidate therapeutic targets, validated genes were ranked using the multi-criteria evidence scoring framework described in the Methods. Genes exhibiting strong differential expression across both discovery datasets, robust validation within the TCGA cohort, enrichment in aggressive molecular subtypes, significant survival associations, and detectable expression within breast cancer cell lines received the highest composite evidence scores (Figure 12).

**Figure 11a.**
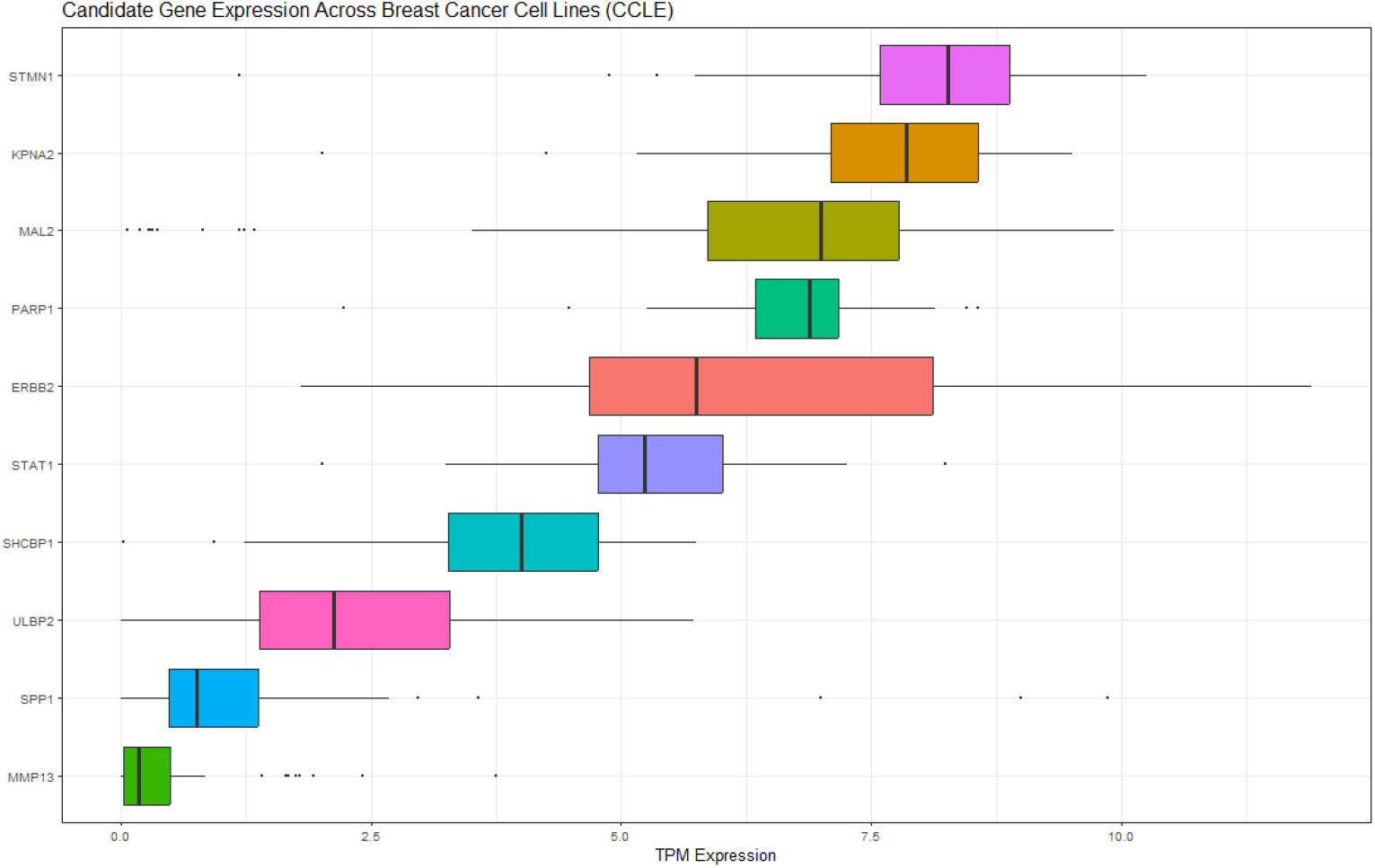
Boxplot of expression levels for candidate genes across breast cancer cell lines from the Cancer Cell Line Encyclopedia (CCLE; 22Q2 release). TPM-normalized expression values are shown. A substantial proportion of candidate genes show detectable expression across cell lines, supporting experimental tractabilitys.

**Figure 12.**
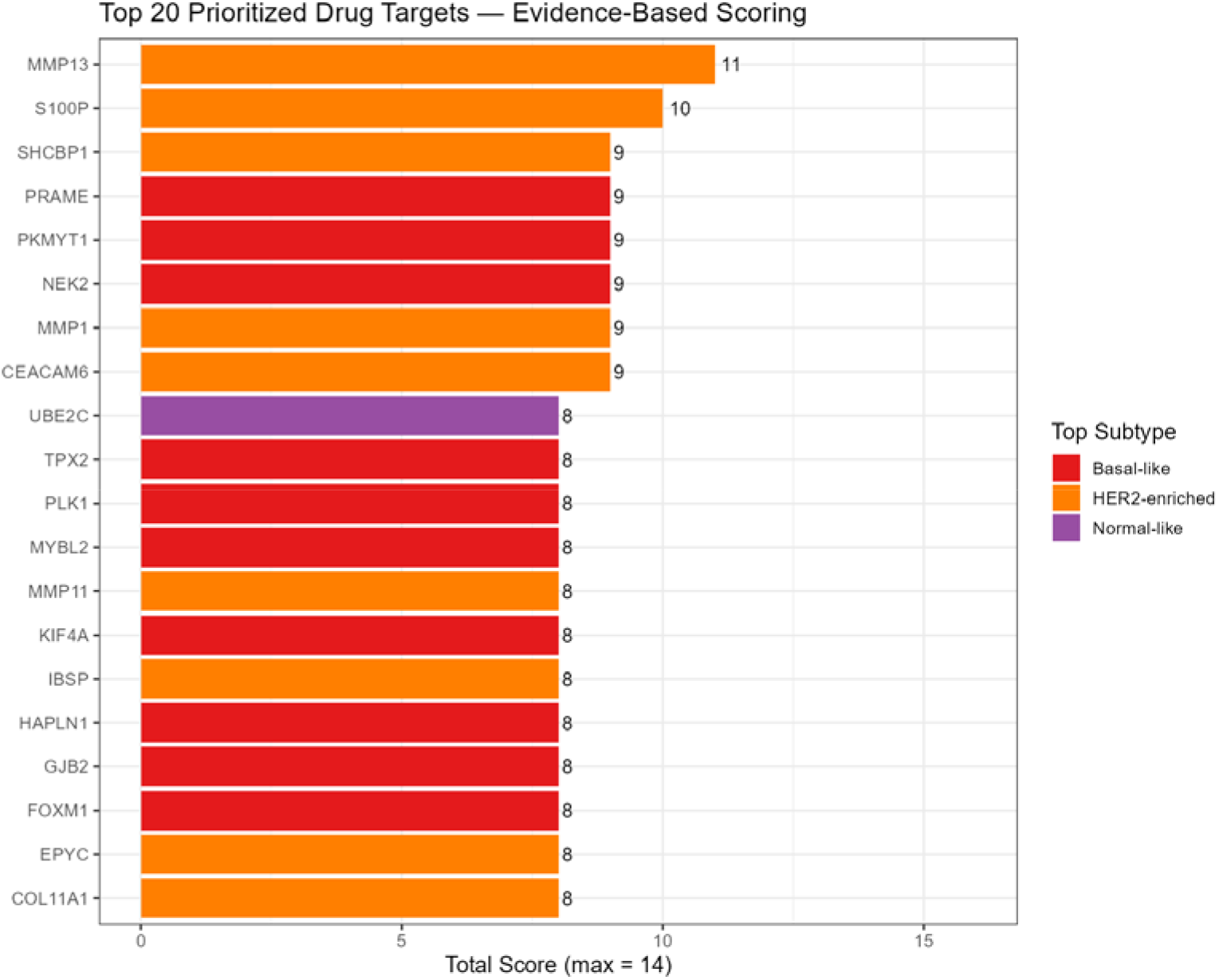
Target prioritization scoring plot for candidate therapeutic genes. Genes are ranked by composite evidence score (maximum 10 points) incorporating differential expression strength, validation robustness, subtype specificity, survival association, and cell line expression. The full scoring matrix is provided in Supplementary Table 1.

Among the highest-ranked candidates were genes associated with proliferative signaling, chromosomal instability, extracellular matrix remodeling, and immune-related pathways. Collectively, these findings demonstrated that integrating independent RNA-seq workflows with downstream clinical and biological validation can systematically identify robust candidate biomarkers and therapeutically relevant targets in breast cancer.

## Discussion

In this study, we developed and applied a three-phase integrative transcriptomic framework to identify reproducible and clinically relevant gene signatures in breast cancer through cross-platform validation and downstream biological characterization. By integrating differential expression analyses from two independently processed RNA-seq datasets — a STAR-counts-based TCGA-BRCA cohort and a Salmon-based SRA-derived dataset — with large-scale validation in the full TCGA-BRCA cohort, we identified a robust set of consensus upregulated genes associated with aggressive breast cancer phenotypes, proliferative signaling, and adverse clinical outcomes.

A major challenge in transcriptomic research is the variability introduced by differences in RNA-seq processing workflows, dataset composition, and analytical strategies. Although numerous studies have reported breast cancer-associated transcriptional signatures, reproducibility across independent datasets remains inconsistent, limiting translational applicability (Ioannidis, 2005). In this study, we address the reproducibility challenge in computational transcriptomics (Peng, 2011) by demonstrating that 666 genes are concordantly upregulated across two independently processed datasets, indicating that core oncogenic drivers exhibit cross-platform stability that persists through variations in library preparation, quantification algorithms, and cohort demographics. Importantly, the subsequent validation of 502 of these genes in the full TCGA-BRCA cohort, corresponding to a validation rate of approximately 93%, further supports the robustness of the identified signatures.

The strong enrichment of cell cycle-associated and chromosomal instability pathways observed throughout the functional enrichment analyses is consistent with established hallmarks of breast tumor progression. Gene Ontology, KEGG, and Reactome analyses collectively highlighted dysregulation of mitotic progression, chromosome segregation, DNA replication, and homologous recombination pathways. These findings align with previous studies demonstrating that proliferative signaling and genomic instability are central drivers of aggressive breast cancer biology, particularly within Basal-like and HER2-enriched tumors (Cancer Genome Atlas Network, 2012). The recurrent enrichment of Fanconi anemia and DNA repair-associated pathways additionally suggests persistent replication stress and impaired genomic maintenance mechanisms within the identified tumor-associated signatures.

Molecular subtype analysis further demonstrated that the validated gene set was disproportionately enriched in Basal-like and HER2-enriched tumors, which together accounted for approximately two-thirds of dominant subtype-associated genes. This observation is biologically significant because these subtypes are frequently associated with increased proliferative activity, genomic instability, poorer prognosis, and reduced therapeutic responsiveness compared with Luminal tumors. Recent transcriptomic and single-cell studies have similarly shown that Basal-like breast cancers exhibit highly dynamic transcriptional programs linked to cell cycle activation, immune modulation, and tumor microenvironment remodeling (Wu et al., 2021; Kim et al., 2024). The predominance of validated genes within these aggressive subtypes suggests that the integrative framework preferentially captured transcriptional programs associated with biologically active tumor progression rather than subtype-neutral expression changes.

Although survival analysis identified only four genes that remained statistically significant following multiple testing correction, this finding should be interpreted within the clinical context of the TCGA-BRCA cohort. TCGA breast cancer datasets are characterised by relatively favourable survival outcomes and low event rates, which substantially reduce statistical power for survival-based analyses; accordingly, the number of genes surviving stringent correction is expected to be small. Nevertheless, the significant associations observed for *MMP13, ULBP2, SHCBP1*, and *MAL2* are noteworthy, and the variation in effect sizes across these four candidates (Table 1) reflects genuine biological heterogeneity in how these genes influence clinical outcomes rather than a methodological weakness. *MMP13*, which showed the most robust survival association (padj = 1.45 × 10□□), encodes a matrix metalloproteinase previously implicated in extracellular matrix remodelling and tumour invasion across multiple cancer types, and its strong prognostic signal in this cohort is consistent with its established role in promoting metastatic dissemination. *SHCBP1* has been associated with mitotic progression and proliferative signalling in multiple malignancies, with reported roles in spindle assembly checkpoint regulation, consistent with the broader cell cycle enrichment signature identified in this study. *MAL2* is a transmembrane protein previously shown to be overexpressed in breast cancer and associated with resistance to trastuzumab, suggesting a role in therapeutic escape and making it a candidate of particular translational interest. *ULBP2* is a ligand for the NKG2D receptor expressed on natural killer cells; its upregulation in tumour tissue likely reflects a mechanism of immune recognition evasion that enables tumour cells to escape NK-cell-mediated cytotoxicity, contributing to disease progression and adverse clinical outcomes. Notably, several other concordantly upregulated genes, including *SPP1* and *STMN1*, did not reach significance after correction in this cohort, underscoring that cross-platform transcriptional concordance and population-scale validation do not automatically confer prognostic relevance, and that survival associations in TCGA must be interpreted conservatively given the cohort’s clinical characteristics.

An important strength of this study is the integration of multiple analytical layers extending beyond conventional differential expression analysis. Rather than relying solely on statistical significance, the study incorporated cross-dataset concordance analysis, large-scale validation, molecular subtype characterization, survival analysis, and cancer cell line expression profiling to prioritize biologically and clinically meaningful candidates. This approach improves confidence that the identified genes represent stable tumor-associated signals rather than dataset-specific artifacts. The additional incorporation of an evidence-based target prioritization framework further enhances the translational relevance of the study by systematically integrating transcriptomic, clinical, and experimental evidence.

The study also demonstrates the value of combining controlled discovery cohorts with large-scale validation datasets. The balanced 30 tumor versus 30 normal TCGA subset provided a controlled environment for initial signal detection with reduced cohort imbalance, while the independently processed Salmon-based dataset enabled technical concordance assessment across RNA-seq workflows. Validation within the full TCGA cohort subsequently established population-level reproducibility. Together, this design allowed the study to address both analytical reproducibility and biological generalizability, two critical challenges in computational oncology.

Despite these strengths, several limitations should be acknowledged. First, the independently processed RNA-seq dataset consisted of only three tumor and three normal samples, which limits statistical power and reduces sensitivity for detecting low-abundance transcriptional changes; however, the purpose of this dataset was not large-scale biomarker discovery but rather independent workflow validation and concordance assessment. Second, the study relied primarily on transcriptomic data and did not incorporate additional molecular layers such as proteomics, epigenomics, or mutational profiling, which could provide further biological insight into candidate gene function and therapeutic relevance. Third, although candidate genes were validated in breast cancer cell lines, experimental functional validation was not performed and will be necessary to confirm mechanistic roles in tumor progression and therapeutic response. Future work addressing these limitations may incorporate single-cell transcriptomics, spatial transcriptomics, or multi-omics integration to further resolve tumor heterogeneity and microenvironmental interactions, as well as machine learning-based prioritization strategies and external validation cohorts to improve predictive robustness.

In conclusion, this study presents a reproducible and clinically oriented three-phase framework for breast cancer transcriptomic analysis through integration of independent RNA-seq workflows, large-scale validation, and downstream biological characterization. The high degree of cross-dataset concordance observed across independent analytical strategies supports the robustness of the identified gene signatures, while subtype, survival, and cell line analyses highlight their biological and translational relevance. Collectively, these findings contribute to ongoing efforts in reproducible cancer transcriptomics and provide a systematic strategy for prioritizing candidate biomarkers and therapeutic targets in breast cancer.

## Data Availability

TCGA-BRCA RNA-seq data and clinical metadata are publicly available through the Genomic Data Commons (GDC) portal (https://portal.gdc.cancer.gov/) and are accessible via the TCGAbiolinks and curatedTCGAData Bioconductor packages. The independent RNA-seq dataset is publicly available through the NCBI Sequence Read Archive (SRA; accession: **SRP631214)**. Breast cancer cell line expression data were obtained from the Cancer Cell Line Encyclopedia (CCLE) via the depmap Bioconductor package (22Q2 release). All analysis code used and results obtained in this study is available at **<u>Githublink</u>**.

## Notes

https://github.com/jfavourite/TCGA-BRCA-Breast-Cancer-Target-Identification-Pipeline

https://github.com/jfavourite/Transcriptomic-Analysis-of-Breast-Cancer

https://github.com/jfavourite/TCGA-BRCA-Differential-Expression

https://www.ncbi.nlm.nih.gov/biosample?LinkName=bioproject_biosample_all&from_uid=1344446

### Competing Interest Statement

The authors have declared no competing interest.

